# Convection - Diffusion model of talking bacteria

**DOI:** 10.1101/101931

**Authors:** Sarangam Majumdar, Subhoshmita Mondal

**Affiliations:** DISIM, University of L’Aquila, Italy; Department of Chemical Engineering, Jadavpur University, Kolkata 700032, India

**Keywords:** Quorum Sensing, Diffusion, Runge-Kutta Methods, Finite element method, Euler methods

## Abstract

Quorum sensing is cell to cell communication process through chemical signals formally known as autoinducers. When the concentration of quorum sensing molecules reached threshold concentration bacteria are in active state or quorum state. In this article, we propose a mathematical model of quorum sensing systems and study this biological system numerically. Moreover, we compare the different numerical scheme with the batch culture of *P.aeruginosa*. We observed a negative diffusion coefficient which plays an important role in the quorum sensing mechanism.

## 1 Introduction

Nature is full of amazing facts and one such fact is the existence of bacterial cells in the human body. There are trillions of human cells that make each of us but surprisingly the reality is that there are ten trillions of bacterial cells residing in and on a human body! So not more than 10% of human cells are present compared to the huge bacterial cells. It would not be foolish if we introduce ourselves as only 10% human but 90% bacteria! [1]

Bacteria are stronger than a few and thus by union are able to overcome obstacles too great for the few [2]. For many years bacteria were considered as autonomous unicellular organism with the capacity for collective behaviour. In the bacterial world each individual cell reproduces by binary fission and strives to outcompete its neighbours, recognition and co-operation between cells may at first appear very unlikely [3]. Apart from that bacteria produce small diffusible chemicals(pheromones) which was originally coined by Karlson and Lu¨scher (1959). This specific signaling molecules sometimes term as autoinducers [4]. As the bacterial culture grows, autoinducers are released into the extracellular milieu and accumulate. When a threshold concentration of the molecule is achieved, a co-ordinated change in bacterial behavior is initiated. This phenomenon is known as quorum sensing (1994) [5]. The size of the quorum is not fixed but will vary according to the relative rates of production and loss of signal molecules. The bacterial cells are integrated in order to determine their optimal survival strategy. Thus quorum sensing is an integral component of the global gene regulatory networks which are responsible for facilitating bacterial adaptation to environmental stress [3].

**Figure 1:**
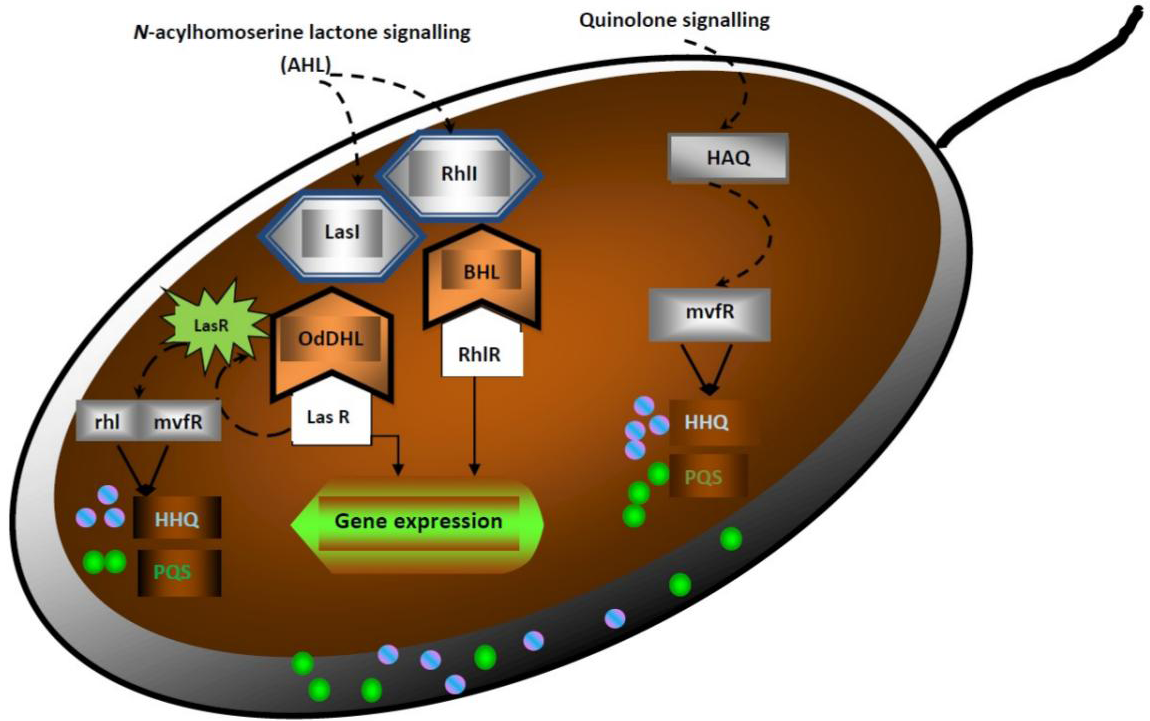
Cascade of bull regulatory network in *Pseudomonas aeruginosa*. *Pseu-domonas aeruginosa*. consist of two LuxR/LuxI-type quorum sensing systems termed as las and rhl. The bull activity is induced by two pathways, (i) N-acylhomoserine lactone signalling (AHL) and (ii) Quinolone signalling (non-AHL signalling). In the first case, LasI and RhlI produce acyl-homoserine lactone autoinducer N-3-oxododecanoyl homoserine lactone (OdDHL) and N-butanoyl homoserine lactone (BHL) respectively. Next, OdDHL and BHL bind to their cognate receptors LasR and RhlR. Activated LasR and RhlR induce expression of a complement of genes, including their own loci. This completes the autoinducing circuits. Also, activated LasR enhances the expression of loci rhl and mvfR. These two loci increase the production of 4-hydroxy-2-heptylquimoline (HHQ) and Pseudomonas quinolone signal (PQS). On the other hand, in quinolone signalling, 4-hydroxy-2-alkylquinolines (HAQ) is controlled by transcriptional receptor mvfR which modulates the expression of genes involved in the production of HHQ and PQS. The las system is placed above rhl and mvfR system in the circuit. Finally, hydrophobic PQS pass to the neighbouring cells through small extra-cellular membrane vescicles.

## 2 Cell-to-cell communication

The quorum sensing field began five decades ago where seminal papers on pheromone-like systems in bacteria highlighted at the intriguing possibility that individual bacterial cells had ambitions beyond dividing into two, in fact, that communication and cooperation were commonplace in the prokaryotic world. McVittie(1962) [6], Tomasz(1965) [7], Khoklov(1967) [8] and Nealson(1970) [9] are the pioneer researchers in this context. They discovered two different types of mediated quorum sensing so far. One is acylhomoserine lactone (AHL) mediated quorum sensing which is very common in gram negative bacteria; another is peptide mediated quorum sensing in gram positive bacteria. This peptide known as autoinducing peptide (AIP) ranges from 5 to 34 amino acids in length and typically contains unusual chemical architectures. Experimentally identified quorum sensing molecules are 4-quinolones, fatty acids and fatty acid methyl esters for Gram negative bacteria. Gram positive bacteria employ unmodified or post-translationally modified peptides. However, we still have not detected any universal quorum sensing systems or quorum sensing molecules. AHL dependent quorum sensing is discovered in Gram negative bacteria like *V.fischeri*, *Pseudomonas aeruginosa* and *Serratia marcescens*.

Quorum sensing is considered in the context of cell cell signaling between intra and inter species. The communication process is coordinated by quorum sensing molecules (autoinducer). QSM have some characteristics feature (1) the production of the quorum sensing signal takes place during specific stages of growth, under certain physiological conditions, or in response to environmental changes; (2) the quorum sensing signal accumulates in the extracellular milieu and is recognized by a specific bacterial receptor; (3) the accumulation of a critical threshold concentration of the quorum sensing signal generates a concerted response and (4) the cellular response extends beyond physiological changes required to metabolize or detoxify the molecule. When all four criteria are met, a molecule can be classified as QSM. The most interesting feature of these quorum sensing signals were well defined in the year 2002 by Winzer et al. These metabolites can induce, during their release, their own uptake system and the production of enzymes required for their breakdown. This indirectly influences the expression of genes from other linked metabolic pathways. [10, 11]

Different kinds of bacteria use different QSM. We have listed a few of them here in the following table and figures.

**Figure 2:**
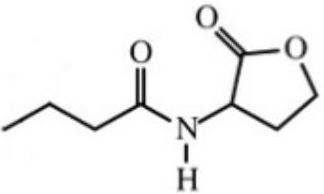
Structure of C4-HSL

**Table 1:** Quorum sensing molecules used by different bacteria

| Signal | Organisms |
| --- | --- |
| C4-HSL (an AHL) | *Aeromonas hydrophila*, *Pseudomonas aeruginosa* |
| C6-HSL | *Erwinia carotovora*, *Pseudomonas aureofaciens*, *Yersinia enterocolitica* |
| 3-Oxo-C6-HSL | *E. carotovora*, *Vibrio fischeri*, *Y. enterocolitica* |
| 3-Oxo-C8-HSL | *Agrobacterium Tumefaciens* |
| Autoinducing Peptide (AIP)-I | *Staphylococcus aureus* Group I strains |
| AI-2 (S-THMF-borate) | *Vibrio harveyi* |
| Competence and Sporulation Stimulating Factor (CSF) | *Bacillus subtilis* |
| Farnesol | *Candida albicans* |

**Figure 3:**
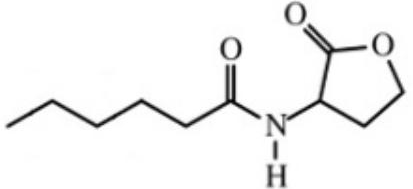
Structure of C6-HSL

**Figure 4:**
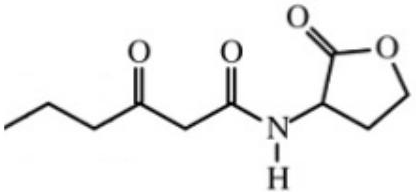
Structure of 3-Oxo-C6-HSL

**Figure 5:**
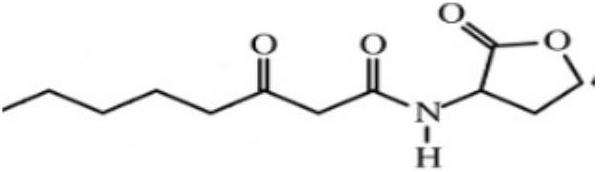
Structure of 3-Oxo-C8-HSL

**Figure 6:**
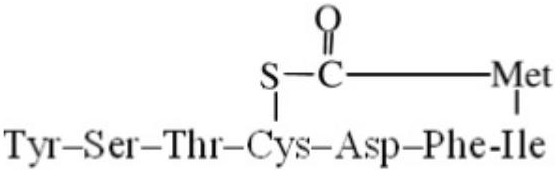
Structure of Autoinducing Peptide (AIP)-I

**Figure 7:**
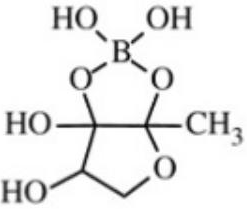
Structure of AI-2 (S-THMF-borate)

**Figure 8:**
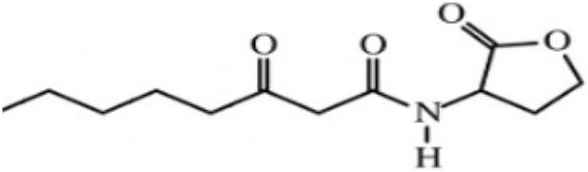
Structure of Farnesol

## 3 Mathematical Model

Mathematical modelling is one of the important tools to study natural systems. Researchers like Anguige, Dockery, Ward and Majumdar have developed several mathematical models of quorum sensing systems based on experiments and analysed the system in detail [12–20]. Still, certain important facts about QS systems are not understood properly. The hydrodynamics and the origin of noise in the QS mechanism is undiscovered. In this present work, we proposed a new mathematical model of this biological system to investigate hydrodynamic conditions and time dependency of the whole system.

### 3.1 The Model

Our modelling approach is to assume a well mixed population of cells, modelling cell growth and QSM production rate as the convection - diffusion equation. For a bacterial population growing as a batch culture, the typical timescale for QSM concentrations to peak, often corresponding to maximal cell densities, is around 8-14 hours for an initial low density inoculum. The modelling assumptions can be summarized as follows [12].

- The concentration of QSM *u* refers here to the external concentration. The QSM of V. fischeri is freely diffusible across the cell membrane. However, this is not the case for all quorum sensing bacteria. For example the primary QSM of *P. aeruginosa* is not freely diffusible, but is biochemically pumped across the cell membrane, leading to differences in internal and external concentrations. The timescale for transport across the cell membrane to equilibrate is only about five minutes, much less than the timescales of interest, and hence we can assume that the internal and external concentration are directly proportional.
- *ρ*(*x*) is consider as mass density.
- *a* is the diffusion coefficient.
- *b* = φε, where φ is assumed as the average velocity of the QSM and ε is known as the porosity (ratio of liquid volume to the volume).

*R* = *cu* is the source of the QSM concentration. *R* > 0 means the quorum sensing is switched on.

Let Ω be a bounded open set in 
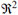
 We consider the space *H* = *L*^2^(Ω) and *V* = *H*_0_^1^ (Ω). We take 
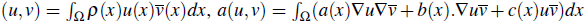
. We make the following hypotheses: ρ and 
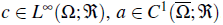
, *divb* = 0, in Ω and 
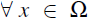
0 < ρ_0_≤ρ(*x*)≤ ρ_M_, 0 < *a*_0_≤*a*(*x*)≤*a*_*M*_, 0≤*c*_0_≤*c*(*x*)≤*c*_*M*_. In the case *V* = *H*^1^(Ω), we will also assume that *c*_0_ > 0 and *b: n* = 0 on the boundary ∂Ω. It is clear that (.,.) defines on *L*^2^(Ω) an inner product equivalent to the usual one and that the sesquilinear form a(.,.) is bounded on *V* × *V*. Besides integration by parts, we have (if *v*ϵ*V*)

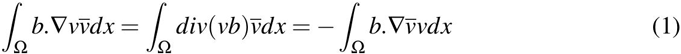

and hence each term of these equations is purely imaginary. Therefore ∀*v* ∊ *V*, *Rea*(*v*, *v*) ≥ 
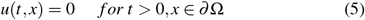
 which prove the ellipticity of *a*.

According to the theory developed in the previous paragraphs, we have a *mα*-accretive operator *A*, and we obtain the mathematical model of the quorum sensing system as follows

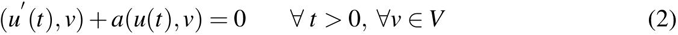

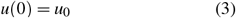

where *u* ∈ *C*^1^((0, ∞);*V*)*C*^0^([0,∞);*H*).The solution of the governing differential equation is unique. Formally (4.2) can be written in terms of a parabolic partial differential equation as follows

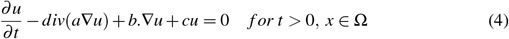

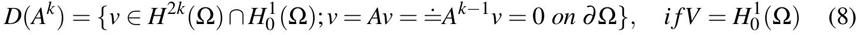

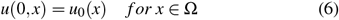

If the initial value u_0_ is real valued, it is clear that the solution *u* is also real valued. Indeed *u* and 
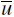
 are both solutions and this solution is unique. On the other hand from the previous analysis provides smoothness results in time *u*∊ *C*^∞^((0; ∞)); *H*) ∩ *C*^*l*^ ([0; ∞); *H*) whenever *u*_0_∊ *D*(*A*^*l*^), *l* ≥ 0. Under this same hypothesis, we also have smoothness in space: *u* ∊ *C*^0^([0; ∞); *D*(*A*^*l*^)) ∩ *C*^*l* − *k*^([0; ∞); *D*(*A*^*k*^)). In order to explicit these results, we can use the regularity theorems for elliptic problems. They give

- Under the regularity assumptions above on the coefficients *a*; *ρ*; *b*; *c* and if, either *ω* in convex, or its boundary is of class *C*^2^, then for *V* = *H*_0_^1^(Ω)

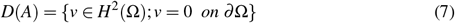

and if 
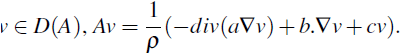

Moreover, if we assume that the boundary of Ω, as well as the coefficient *a, ρ, b, c* are of class *C*^∞^, then for all integer *k* ≥ 1, we have

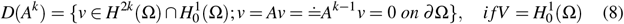

## 4 Experimental Work

The assumption of the well stirred population of bacteria and uniform QSM concentration is well suited to bacterial growth in batch cultures whereby a colony is grown from a low initial inoculum in a flask containing a suitable liquid growth median which is incubated in a cabinate at a constant temperature on a stage that rotates or vibrates. The experiment, specifically designed to give suitable parameter estimates, focuses on growing several strains of *P.aeruginosa* in various media. The cultures were grown for up to 24 hours at 37 degree centigrade and at regular intervals, samples were taken to measure bacteria densities and the QSM concentrations. QSM concentrations were determined using the E. coli bioluminescent reporter strain (*pSB*1142), whereby the bioluminescence from the reporter strain grown in the supernatant from a batch culture was compared with that of media containing a known concentration of (synthetic) QSMs. Bacterial growth in batch cultures generally follows a standard pattern: usually there is an initial lag phase (1-2 hours) followed by a 6-12 hours phase of exponential growth until a limiting population density is reached, after which there is phase in which the density remains roughly constant (stationary phase). It is well known that the environmental stresses induced during the stationary phase lead to numerous changes in bacterial behaviour, including perhaps QSM production; however, we note that typically the population becomes quorate during the exponential phase of growth [9,12,21].

## 5 Discussion

In this section, we are analysing the quorum sensing system with the proposed model. We are using different numerical approaches such as Euler methods, Runge-Kutta methods, Finite difference approaches, Finite element methods. We analyse and compare the numerical simulations. Moreover, we study the experimental approach of the system.

### 5.1 Numerical Euler and Runge-Kutta Scheme

Euler methods (explicit, implicit, modified) are the most basic numerical integrations of the ordinary differential equations. This is the first order method, which means that the local error is proportional to the square of the step size and the global error is proportional to the step size. Figure 4.9, Figure 4.10, Figure 4.11 and Figure 4.13 illustrate the quorum sensing molecule concentrations with time. In the initial lag phase (1 - 2 hours) followed by a 6 - 12 hours phase of exponential growth of the population density. Numerical solutions of the proposed model by the Euler forward scheme, Euler backward scheme, and modified Euler scheme also show the exponential growth of the populations. The Euler forward method gives less error than the Euler backward method. The modified Euler solution is more accurate than others.

**Figure 9:**
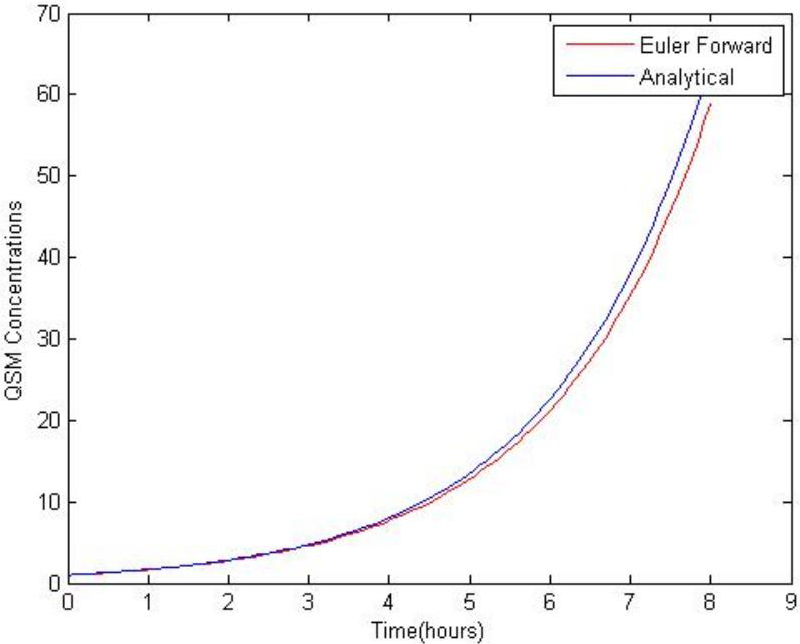
QSM concentration graph by Euler-Forward Method

**Figure 10:**
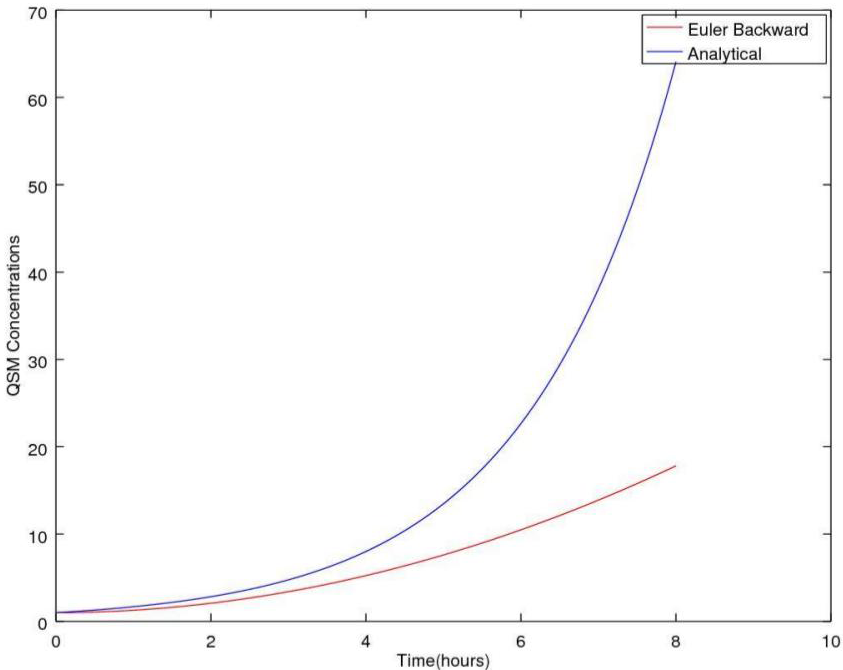
QSM concentration graph by Euler-Backward Method

**Figure 11:**
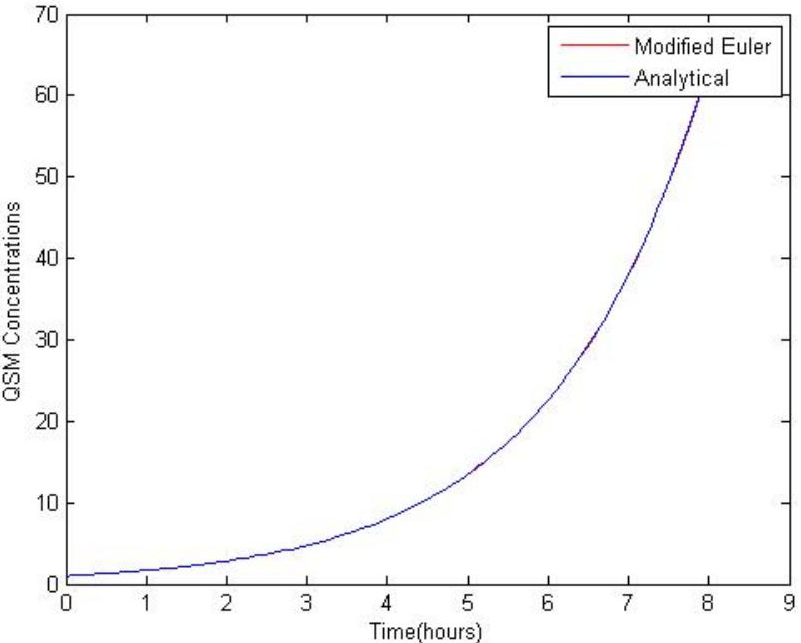
QSM concentration graph by Modified-Euler Method

An important observation is that we get a negative diffusion coefficient in the proposed model which shows that anti diffusion play an important role in the quorum sensing system. Due to the negative diffusion coefficient the bacterial population density continuously rises to a certain level. The diffusible quorum sensing molecules are released, then they accumulate and increase the population density until the threshold concentration is reached. If all the molecules are diffused then quorum sensing never happens. So anti diffusion is an important cause of quorum sensing. The chemical diffusion occurs in the presence of the concentration (or chemical potential) gradient and it results in the net transport of mass. This is the process described by the diffusion equation. This diffusion is always a non-equilibrium process, increases the system entropy, and brings the system closer to equilibrium. The thermodynamic factor is related to the Gibbs free energy. A negative diffusion coefficient means that the flux of *u* diffuses up against the concentration gradient (though still along the chemical potential or free energy gradient). This is known as uphill diffusion, which is important for a special phase transformation, called Spinodal Decomposition.

We use Runge-Kutta explicit and implicit schemes. Figure 4.12, Figure 4.14 shows the concentration graph over time. In this case, we also find the exponential curve in the lag phase. Runge-Kutta order 4 gives a more accurate result than the Runge-Kutta order 2. Both second and forth order methods are convergent and stable. Figure 4.15 illustrates the stability domain of the Euler method, Runge-Kutta method of order 2 and Runge-Kutta method of order 4. The stability of the methods can be interpreted as numerical solutions by the famous Dahlquist test equation. The set 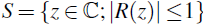 is called the stability domain of the method. If the Runge-Kutta method is of order *p* then, 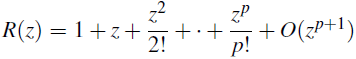. The implicit Runge-Kutta method is also L-stable because it is A-stable and 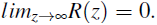 In our case, all the partial derivatives of the function of order *p* exist (and continuous), so that the local error admits the rigorous bound.

**Figure 12:**
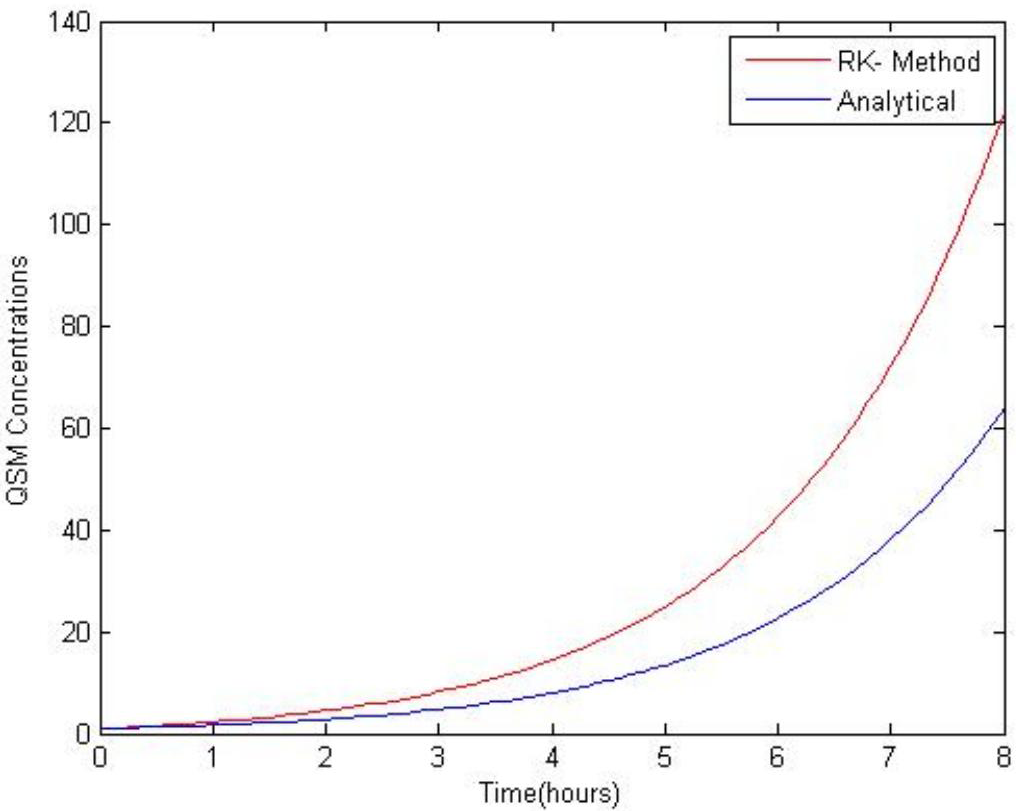
QSM concentration graph by Runge-Kutta Method

**Figure 13:**
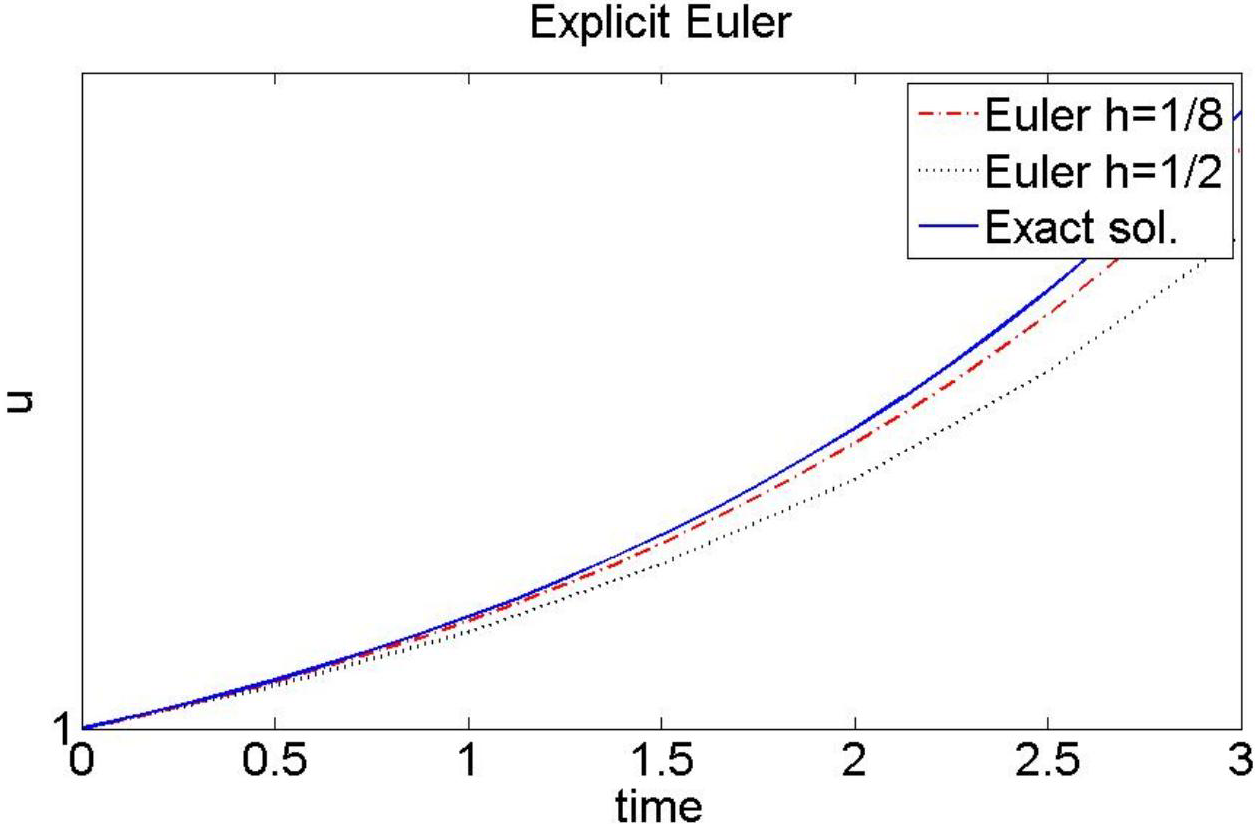
QSM concentration graph by Euler Method

**Figure 14:**
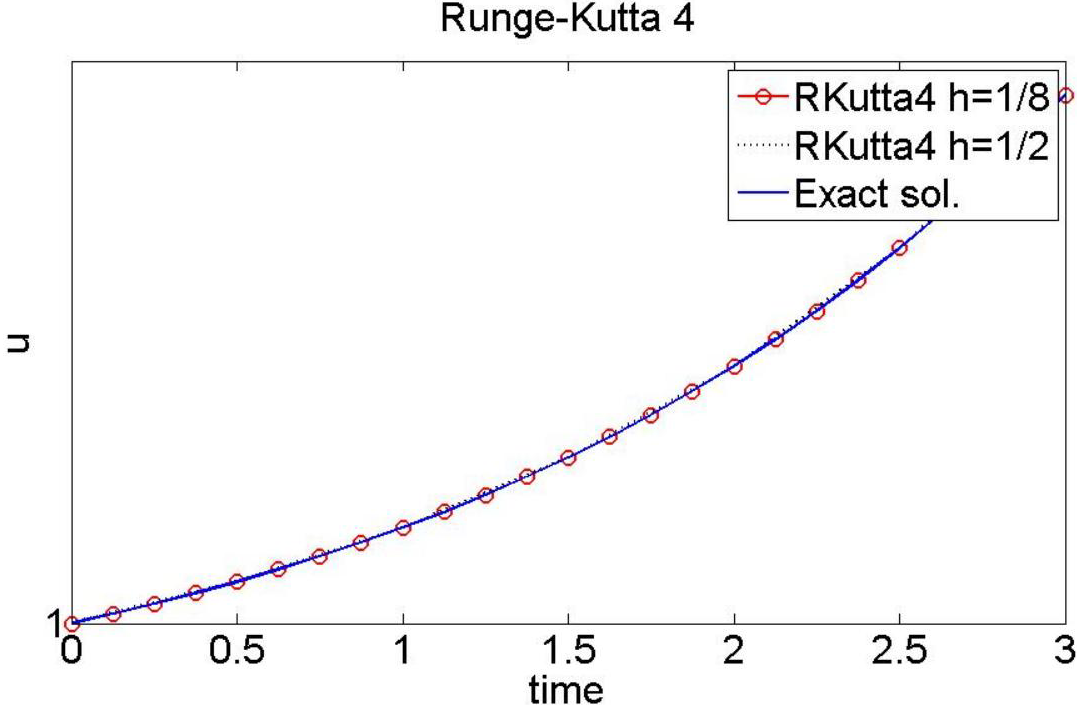
QSM concentration graph by Runge-Kutta 4 Method

**Figure 15:**
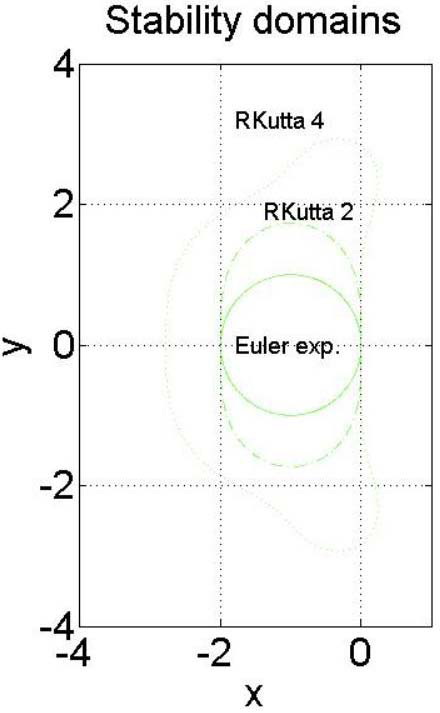
Stability domain

### 5.2 Finite Element Analysis

In this section, we analyse the proposed model of the quorum sensing mechanism using numerical approximation techniques for differential equations which is formally known as finite element approximation. We discretize the whole domain and analyse the concentration behaviour in space. Figure 4.16 illustrate the QSM concentration in space for the 10 nodes using FEM. Figure 4.17 gives the log(error) vs log(N) graph of the system. Moreover, the finite element discretization gives more accurate results than other previously discussed methods.

**Figure 16:**
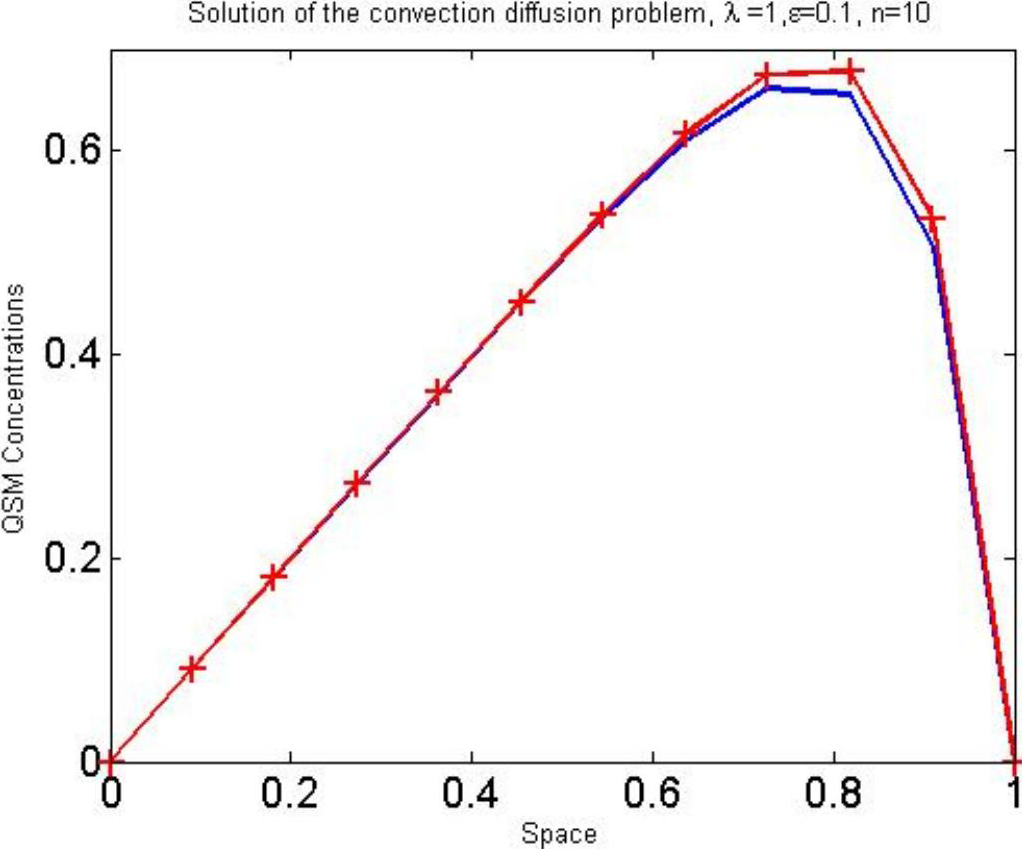
Concentration graph in space using Finite element method

**Figure 17:**
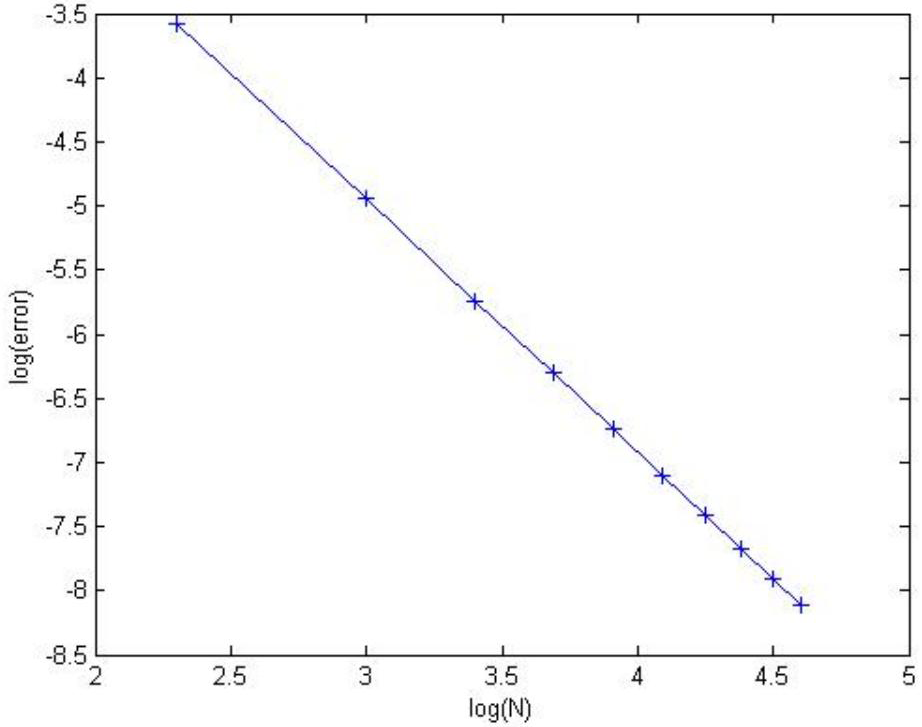
Log(N) vs log(error)

Figure 4.18, Figure 4.19, Figure 4.20 and Figure 4.21 shows the QSM concentration behaviour for different discretizations (n =10),(n=100),(n=1000) and (n=10000) respectively using finite element methods. In every case the most common observation is that the concentration is increased in space because of talking bacteria.

**Figure 18:**
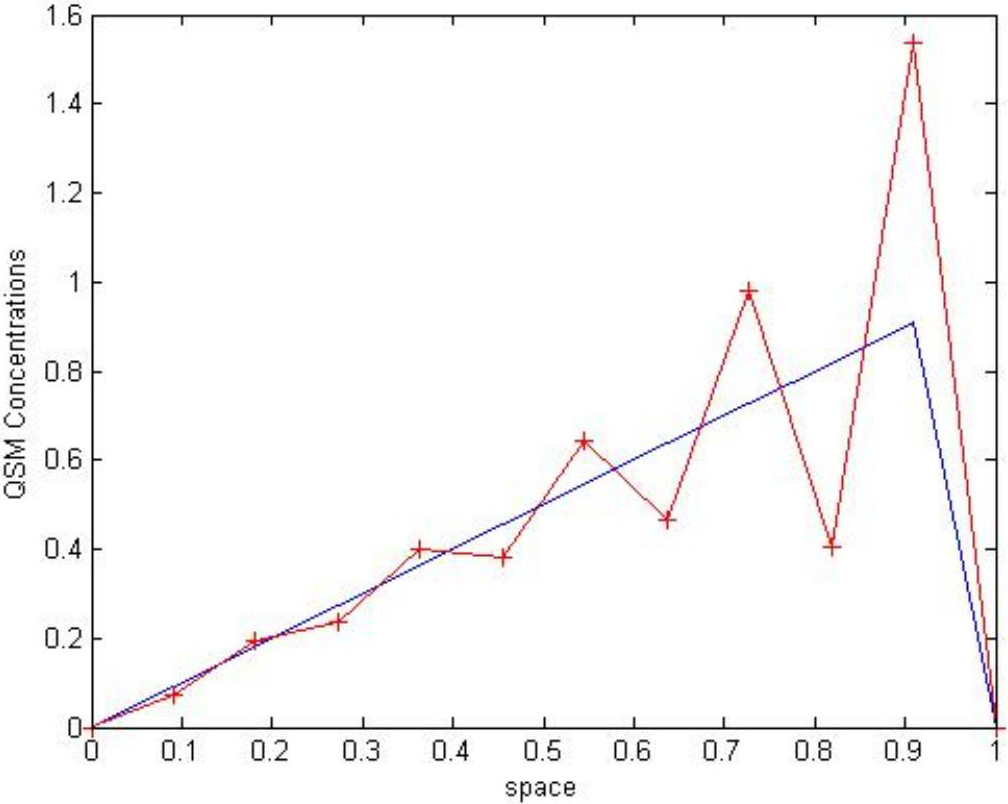
(For n=10) concentration behaviour using FEM

**Figure 19:**
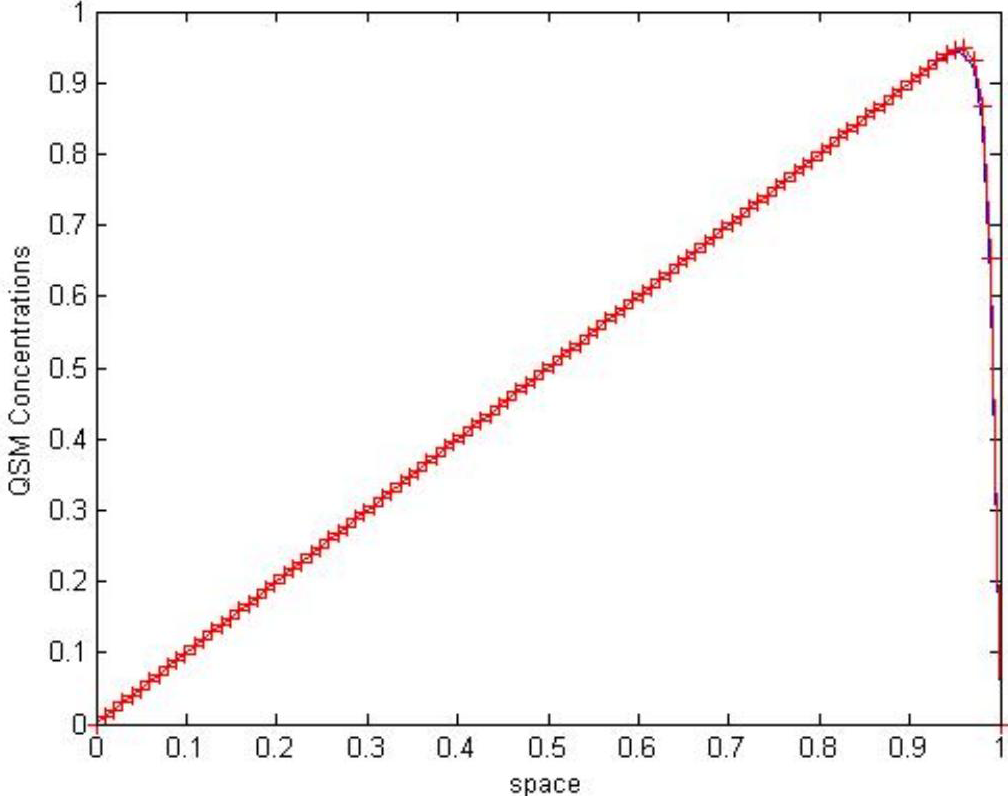
(For n=100)concentration behaviour using FEM

**Figure 20:**
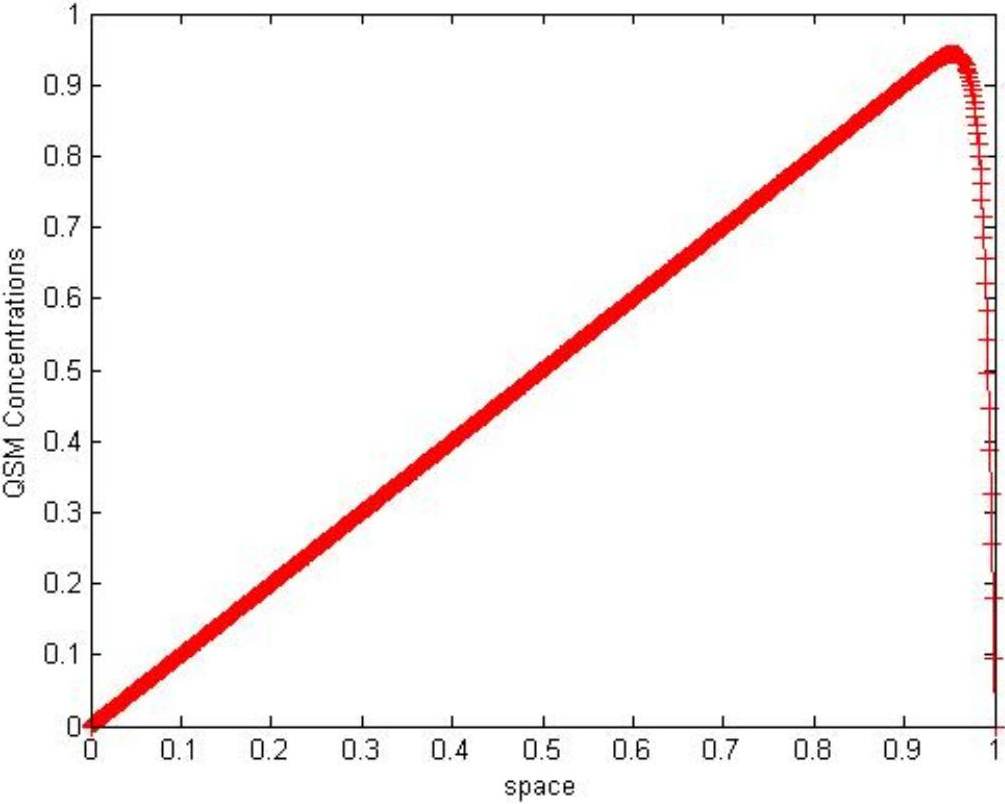
(For n=1000)concentration behaviour using FEM

**Table 2:**
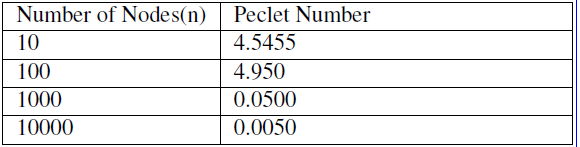
Peclet numbers with different nodes

**Figure 21:**
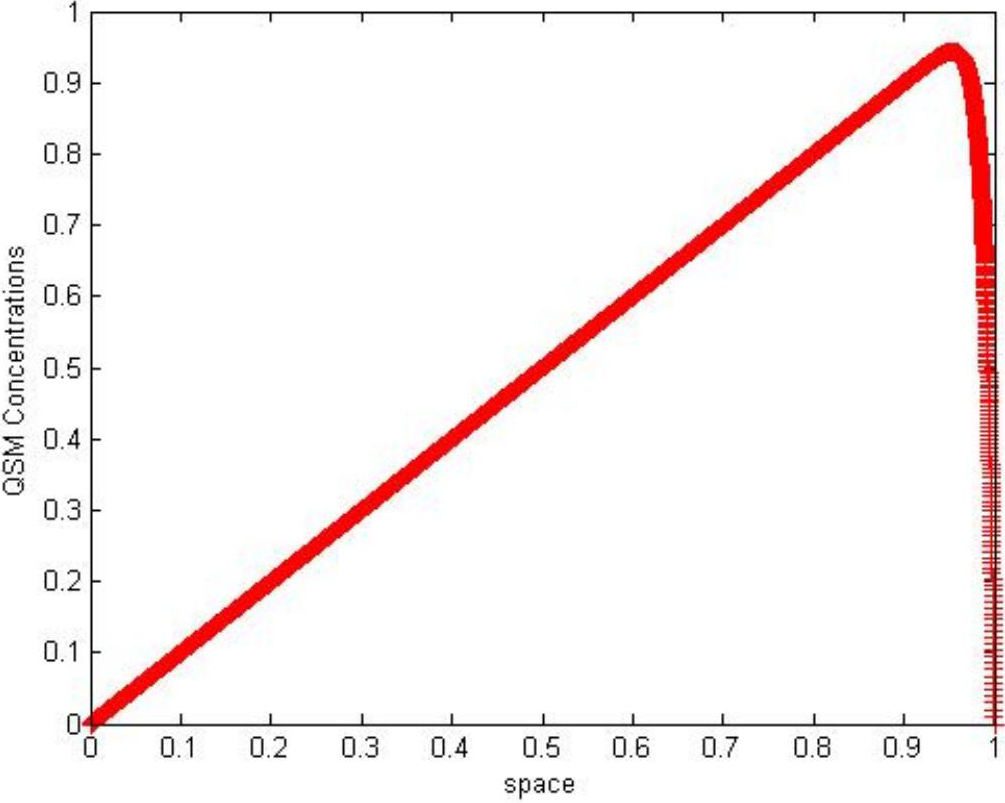
(For n=10000)concentration behaviour using FEM

Peclet number is a dimensionless independent mass transfer parameter which is defined as the ratio of the advective transport rate and diffusive transport rate. For mass transfer, we can write *Pecletnumber* = *Reynoldsnumber* × *Schmidtnumber*. Table 4.2 illustrates the Peclet number for different number of nodes.

### 5.3 Numerical Experiment

We are now showing some experimental evidence of the behaviour of the QSM concentration in time and space. We also compare the numerical result with the experiment data in this section. Figure 4.22, Figure 4.23 show the numerical experimental graph of the concentration with time. We can easily say that concentration is increased in time and reaches a threshold level before falling down.

**Figure 22:**
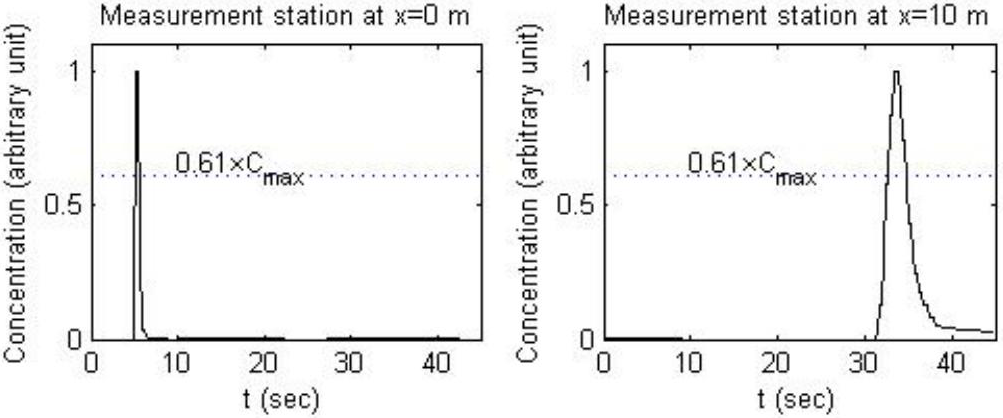
Concentration(Numerical experiment)graph

**Figure 23:**
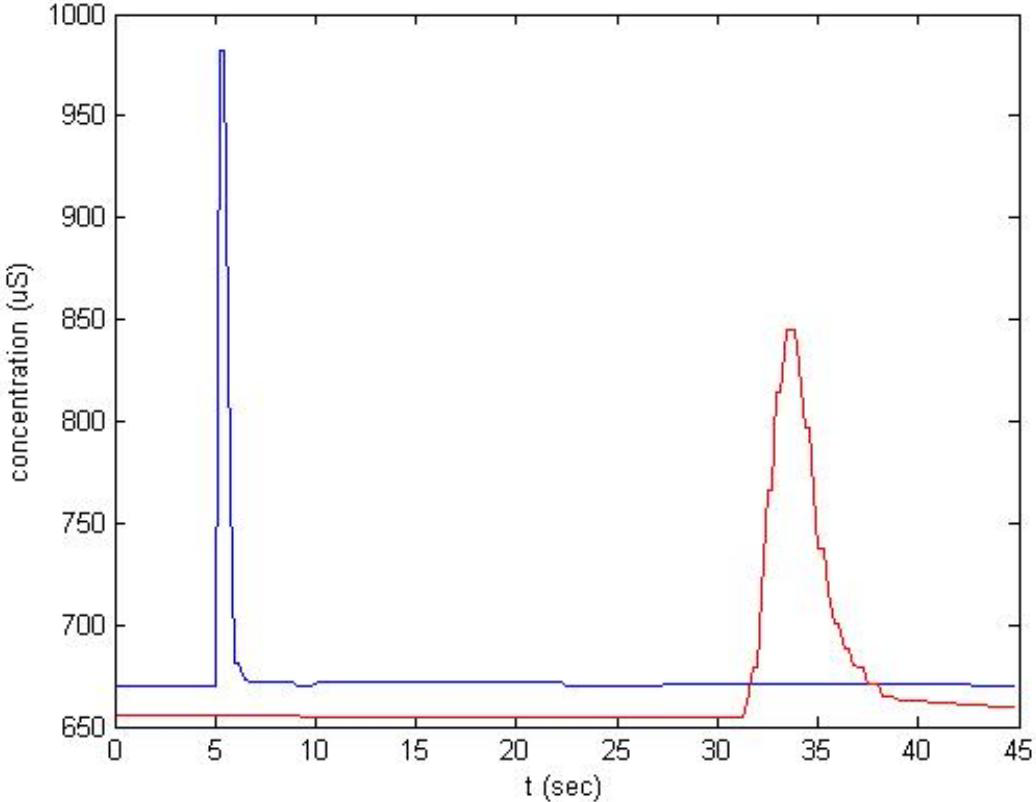
Concentration graph of experiment

The following graph shows the comparison of the experiment data and the numerical result of the concentration behaviour for only the convection effect at different times. Figure 4.24, Figure 4.25, Figure 4.26, Figure 4.27 and Figure 4.28 illustrate concentration behaviours at (time = 0.05), (time = 0.1), (time = 0.15), (time = 0.2) and (time = 0.25) respectively.

**Figure 24:**
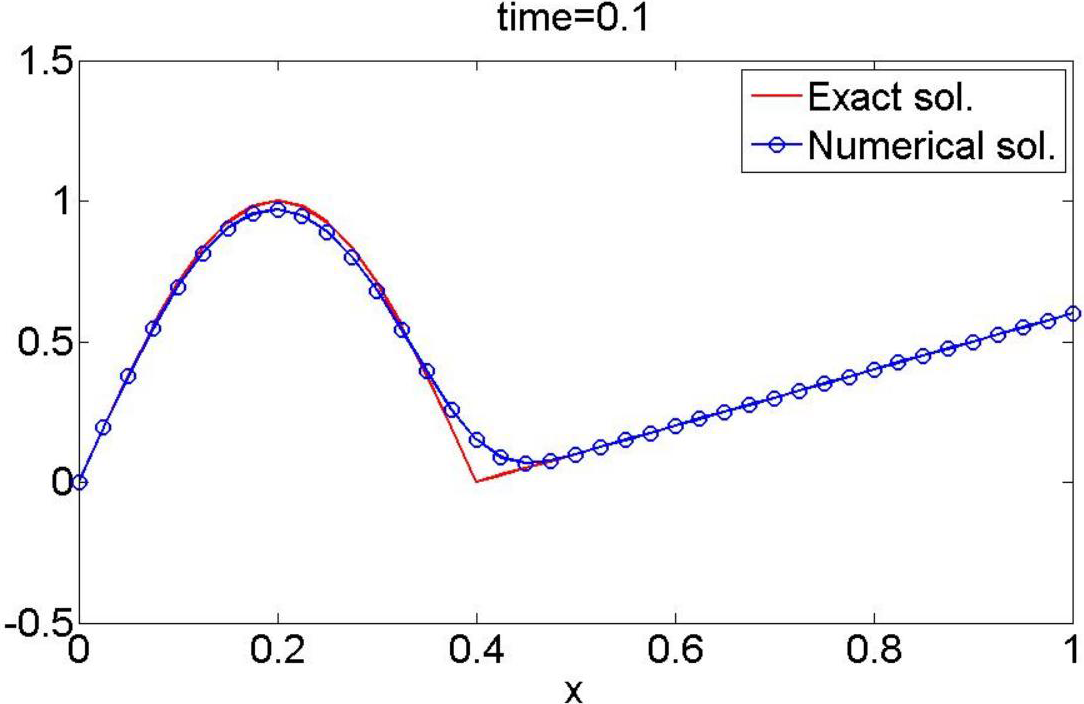
Concentration behaviour graph for only convection effect at time=0.05

**Figure 25:**
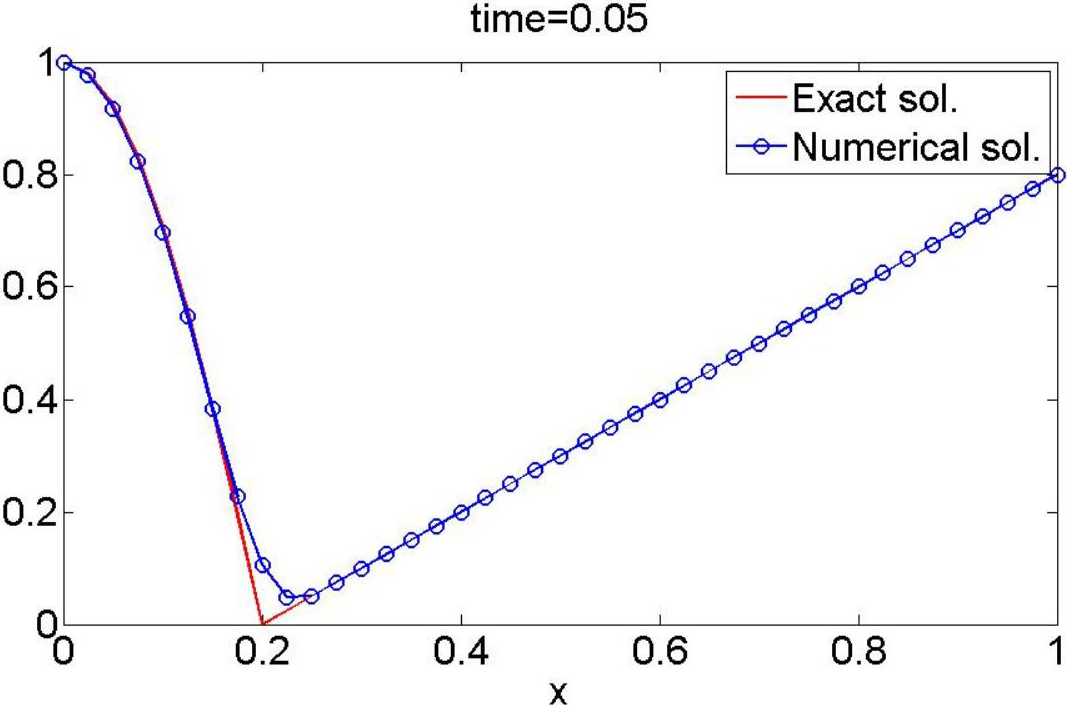
Concentration behaviour graph for only convection effect at time=0.1

**Figure 26:**
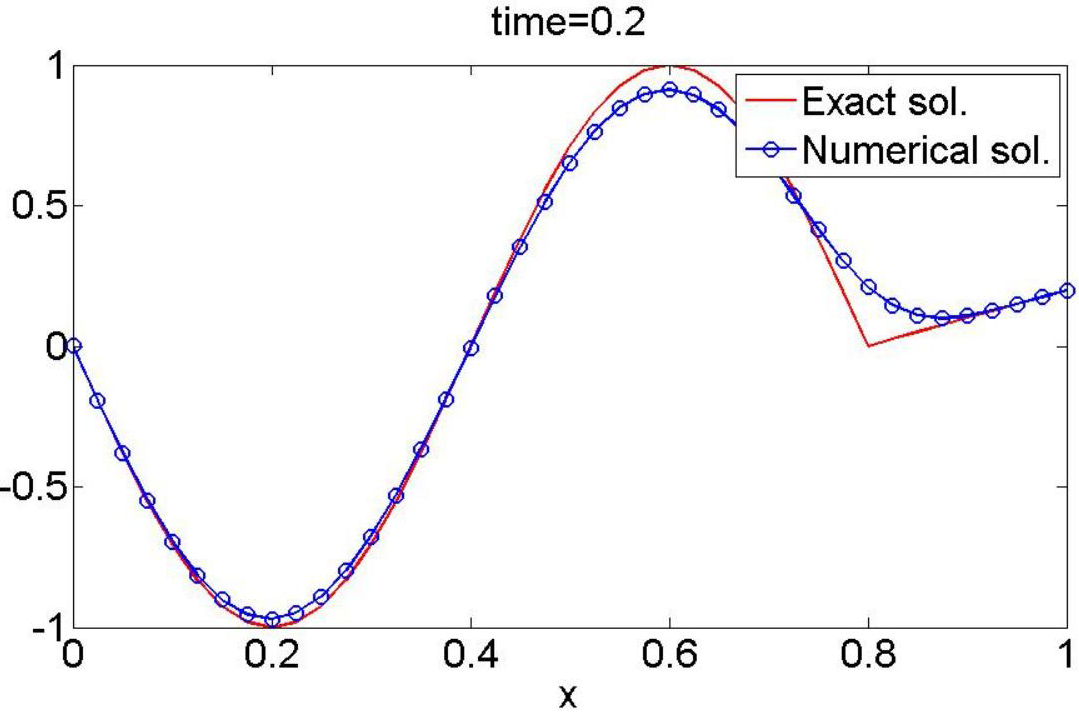
Concentration behaviour graph for only convection effect at time=0.15

**Figure 27:**
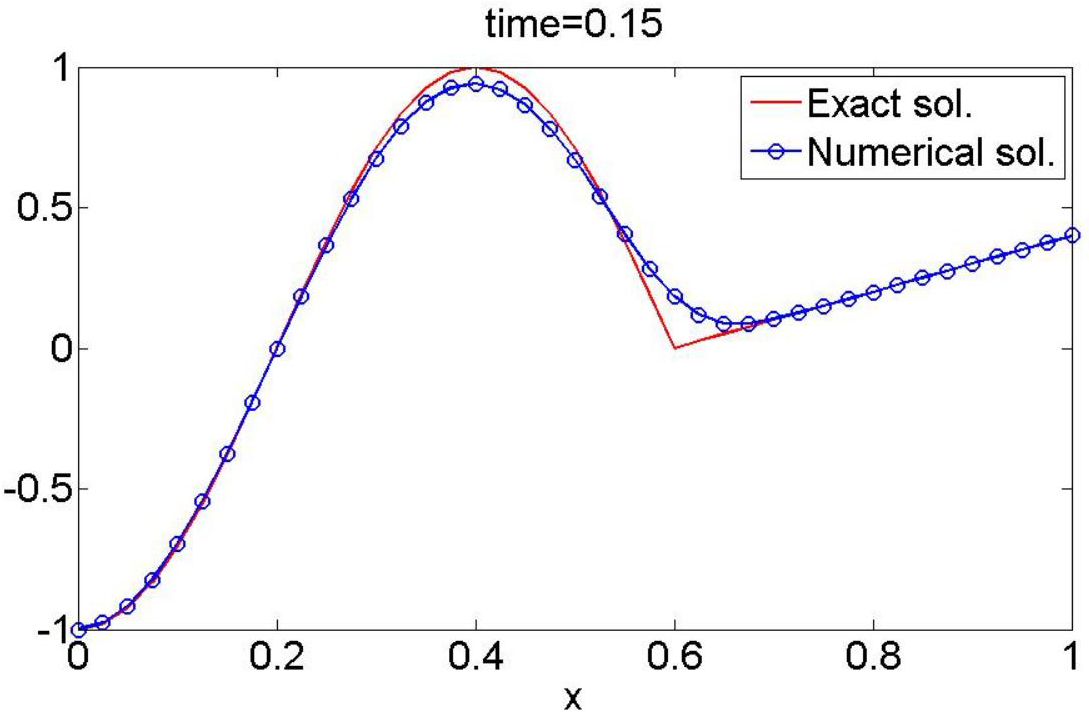
Concentration behaviour graph for only convection effect at time=0.2

**Figure 28:**
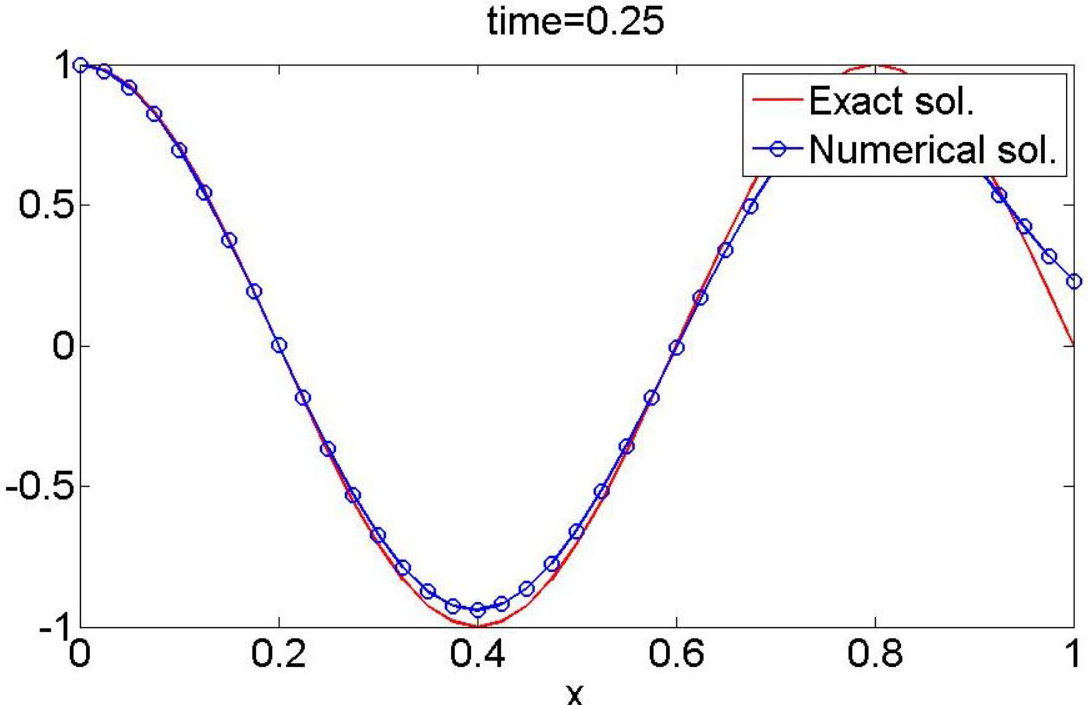
Concentration behaviour graph for only convection effect at time=0.25

## 6 Conclusion

We applied this mathematically sound approach to investigate the quorum sensing system in bacteria. We proposed a mathematical model of the quorum sensing mechanism which is basically a convection - diffusion model. The model predicted the behaviour of the QSM concentration behaviour in space and time. We observed a negative diffusion coefficient which occurred in this complex biochemical phenomenon. We used various numerical schemes (explicit Euler, Implicit Euler, explicit Runge-Kutta, implicit Runge-Kutta, Finite element method) and compared the results in the sense of the approximation of the solution and the stability of methods. We observed that the finite element approximation gave a better result than others. Moreover, the Runge-Kutta scheme also gave A-stability and concentration curves in time which are more accurate with the experimental data. The batch culture of the quorum sensing system and our proposed system give approximately good results. Finally, we did some numerical experiments with this density dependent behaviour of the bacterial talk with only convection effect which illustrated a behaviour of the quorum sensing in space and time.

